# Field application of de novo transcriptomic analysis to evaluate the effects of sublethal freshwater salinization on *Gasterosteus aculeatus* in urban streams

**DOI:** 10.1101/2023.08.14.553225

**Authors:** Camilo Escobar-Sierra, Kathrin P. Lampert

## Abstract

Freshwater salinization poses global challenges for aquatic organisms, impacting their physiology and ecology. However, current salinization research predominantly focuses on mortality endpoints in limited model species, overlooking the sublethal effects on a broader spectrum of organisms and the exploration of adaptive mechanisms and pathways under natural field conditions. To address these gaps, we conducted high-throughput sequencing transcriptomic analysis on the gill tissue of the euryhaline fish *Gasterosteus aculeatus*, investigating its molecular response to salinity stress in the highly urbanized river Boye, Germany. We found that even sublethal concentrations of chloride led to the activation of the energetically costly osmoregulatory system in *G. aculeatus*, evidenced by the differential expression of genes related to osmoregulation. Our enrichment analysis revealed differentially expressed genes (DEGs) related to transmembrane transport and regulation of transport and other osmoregulation pathways, which aligns with the crucial role of these pathways in maintaining biological homeostasis. Notably, we identified candidate genes involved in increased osmoregulatory activity under salinity stress, including those responsible for moving ions across membranes: ion channels, ion pumps, and ion transporters. Particularly, genes from the solute carrier family SLC, aquaporin *AQP1*, chloride channel *CLC7*, ATP-binding cassette transporter *ABCE1*, and ATPases member ATAD2 exhibited prominent differential expression. These findings provide insights into the molecular mechanisms underlying the adaptive response of euryhaline fish to salinity stress and have implications for their conservation and management in the face of freshwater salinization.

## Introduction

The salinization of freshwater is a pressing issue that is garnering increasing attention from researchers and policymakers alike. In recent years, several studies have highlighted the gravity of the problem [1–3]. While freshwater salinization is often solely associated with increased conductivity and total dissolved solids, it is the accumulation of various ions such as Na+, Ca2+, Mg2+, K+, NO3-, SO4 2-, Br, Cl-, among others [3,4]. A variety of factors can lead to heightened levels of ion accumulations, including background concentrations in watersheds, as well as human-driven changes in land use, sea level rise, agriculture, road salts, wastewater, and mining [5]. Given this background, the Ruhrgebiet region in Western Germany offers an excellent case study for examining freshwater salinization. This area is one of the most densely populated in Europe [6] and encompasses a wide variety of activities that are known to contribute to a higher vulnerability of freshwater systems to salinization. Mining is one of the major threats in this region, with coal extraction requiring the infiltration of groundwater that is subsequently pumped into nearby rivers and streams [7]. As a result, mining water discharge has been recorded to reach chloride concentrations as high as 3500 mg/l in the River Lippe which is 175 times higher than the typical natural background chloride concentrations in the world surface freshwater ecosystems of <20 mg/l [8]. However, the implementation of new management measures and the closure of many mines has led to a significant reduction in concentration, with current levels now below 400 mg/l [7]. Despite this progress, the ongoing inputs from abandoned coal mines in Western Germany continue to contribute to freshwater salinization in the region [9]. Another significant factor contributing to freshwater salinization in the Ruhrgebiet region is the changing land cover. As urbanization has intensified, rivers and wetlands have become increasingly surrounded by impervious man-made surfaces. This has led to a greater runoff of road salts and sewage waters, particularly in urban surface waters which turns them particularly vulnerable to salinization [10].

Most freshwater animals are strictly adapted to low ion concentrations in their surrounding water compared to their internal ion concentrations (stenohaline), and spend a lot of energy for osmoregulation by actively pumping ions inside their bodies and excreting water [11]. Thus, hyperosmotic stress due to freshwater salinization poses a significant physiological challenge to many freshwater animals, including fish which must maintain their ion homeostasis in the face of ion loss through diffusion and water absorption across their permeable membranes [12]. Active membrane transport of ions against concentration gradients is required to maintain internal salt concentrations, and increased environmental salinity above the internal ion concentrations leads to greater energy expenditure for osmoregulation [13]. As a result of the advancement in fish osmoregulation physiology research, and their importance in bio-indication, fisheries, and the economy, fish have become essential models for studying the effects of environmental change [14]. Chloride is widely recognized as the most important anion for osmoregulation in freshwater, making it one of the most commonly used indicators to measure salinity effects on fish [4]. The annual upper chloride mean threshold concentrations in German surface waters, as per the European water directive, range between 40-90 mg/L, with Canada and the U.S. having chronic thresholds of 160 mg/L and 230 mg/L, respectively. [8,15]. However, these thresholds have been based on ecotoxicological tests on a limited number of species that often rely on measuring the salinity effect on endpoint survival and may overlook sublethal effects on non-model species. In ecotoxicology, sublethal effects are defined as those that do not directly cause the death of an individual but rather have an effect on individual fitness (i.e. behavior, cognition, physiology) [16,17]. This is critical, as it is recognized that one of the major challenges of understanding the impact of environmental change is determining the point at which sublethal responses to stressors begin to adversely impact the organism [18]. Focusing solely on tissue, organ, or endpoint mortality effects and using a limited number of model species may overlook critical information regarding the physiological impacts of sublethal salinity stress on organisms. Moreover, until recently, studies on fish physiology have relied on technologies with reduced resolution and a limited range of species models [4]. Thus, there is a pressing need to apply novel technologies and widen the model species pool to gain a better understanding of the effects of salinity stress on freshwater species.

In this context, the application of molecular techniques has greatly improved our understanding of the toxicity mechanisms of major anthropogenic ions in various fish species at high resolution. This development aligns with the recent scientific consensus on the analysis of the response of fish to environmental stressors at different levels, including molecular gene expression and potential adaptation [19]. Furthermore, the molecular adaptation to salinity by freshwater species has been identified as one of the major gaps in the research agenda of global freshwater salinization, and transcriptomic technologies have been proposed as a promising approach to tackle it [2]. Transcriptomics allow capturing a snapshot in time of the total gene expression (mRNA) present in a cell under a given condition [20]. And as noted by [21], can reveal sublethal stress thresholds that extend beyond the negative impacts on individual and population fitness. This provides a more comprehensive understanding of species’ habitat requirements, which in turn facilitates their management. Recently, transcriptomics has been applied to identify the function of novel and conserved genes and the underlying pathways of salinity stress in fish species such as *Lateolabrax maculatus*, *Poecilia reticulate*, *Pseudopleuronectes yokohamae*, *Lates calcarifer*, *Acipenser baeri*, *Oreochromis mossambicus* female × *O. urolepis hornorum* male, *Oncorhynchus keta*, and *Gasterosteus aculeatus* [22–29]. While the studies previously cited have provided insight into the mechanisms underlying tolerance thresholds and adaptations to osmotic stress, they have been limited by their reliance on laboratory manipulations and exposure to salinity concentrations that exceed sublethal thresholds. As highlighted by Whitehead & Crawford (2006) [30], it is crucial to investigate expression variation in ecological contexts to fully comprehend the role of gene expression in the adaptive response to environmental stressors. Under natural conditions, organisms are exposed to a variety of fluctuating environmental signals, providing a valuable opportunity to discern gene expression patterns that are not discernible under laboratory conditions [31].

The threespine stickleback (*Gasterosteus aculeatus*) is an opportunistic species that has demonstrated an exceptional capacity for adaptation to diverse aquatic habitats and is widely distributed in the Ruhrgebiet region in Western Germany. One key trait that has enabled this species to thrive in various environments and colonize systems throughout the Northern Hemisphere is its ability to regulate its osmotic balance in response to changes in salinity (Euryhalinity) [32]. Even landlocked freshwater populations of stickleback, which have been separated from the marine environment for thousands of years, possess the osmoregulatory machinery needed to handle abrupt changes in salinity [33]. This suggests that sticklebacks have maintained their physiological adaptation to saltwater despite repeated colonization of freshwater habitats. Their wide distribution and adaptability have made them an important model in the study of adaptive evolution and have been subject to the development of large-scale genetic and genomic resources [34]. Furthermore, experimental evidence has shown that their gill transcriptome responds to subtle changes in environmental salinities within the freshwater range [24]. Although their euryhalinity suggests they can tolerate the salinity changes, it has been noted that the activation of the hyperosmotic osmoregulatory mechanisms comes at a high energetical cost [35]. Cellular remodeling, enzyme expression, and transport protein synthesis linked to salinity changes have been estimated to account for 20-68% of the total energy expenditure in certain fish species [36]. These costs may drive the need for physiological and behavioral optimization to minimize energy expenditure, resulting in narrow physiological tolerance windows and behavioral avoidance of salinities outside the optimal range [35,37]. However, there is still a lack of knowledge on the molecular mechanisms underlying this species’ responses to salinity stress. Therefore, the threespine stickleback provides an excellent model for studying the mechanisms underlying the osmoregulation of a euryhaline opportunistic species under sublethal salinity concentrations.

Taking into consideration the research gaps and opportunities introduced above, we designed a study to understand the physiological effects on *G. aculeatus* under subtle changes in concentrations of chloride from anthropogenic sources in the field. We hypothesize that the salinity gradient in the urban stream will affect the gene expression profile of *G. aculeatus* gills, with an enrichment of osmoregulation-related pathways. We sampled fish from an urban stream in North Western Germany with a historical salinization influence at three stations along a salinity gradient to test this hypothesis. We then evaluated the transcriptomic fingerprint of *G. aculeatus* gills, performed a comparative transcriptomic study, evaluated the gene ontology of the differentially expressed genes and identified candidate genes related to osmoregulation. This research aims to use de novo transcriptomic analysis to evaluate the effects of sublethal freshwater salinization on *Gasterosteus aculeatus* in urban streams, with a focus on identifying molecular adaptation mechanisms and expanding the model species pool to improve our understanding of the sublethal effects of salinity stress on freshwater species.

## Material and methods

### Sampling design

We choose three stations in a gradient of chloride concentration in two tributaries and in the main channel of the river Boye (Figure 1). The station with the highest salinity of the stations is in the Kirchschemmsbach, a stream that has historically affectation by salinity intrusion (Station C). Followed by the main channel of the Boye in its confluence with the Kirchschemmsbach (Station A). And the last one is a station in a stream close to the other stations with no historical or present influence of secondary salinization, which was used as the control site (Station B). The Chloride concentrations in the period of sampling for station B were 12.912 mg/l, Station A 26.378 mg/l, and station C 32 mg/l. In the sites that we sampled, the concentration was below the annual chloride mean upper threshold from German streams in the central highlands which is in the range of 40-50 mg/l [9]. And below the reported chronic or acute toxicity level for species such as trout and fathead minnow [38]. However, both stations B and C are above the typical Cl− concentrations from natural sources in surface freshwater ecosystems that are in general below 20 mg/L [8].

**Figure 1.**
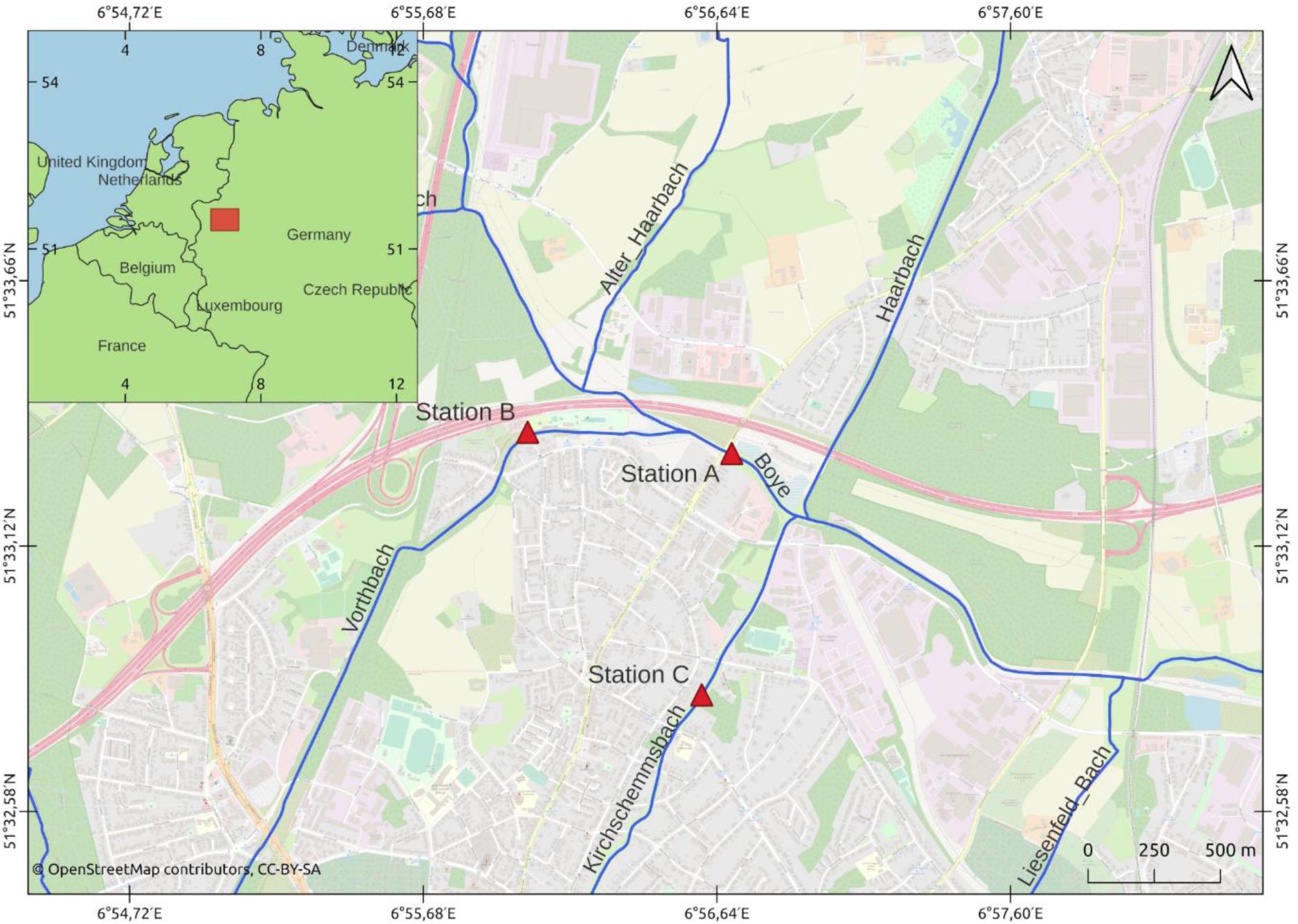
Location of the sampling station in the Boye catchment. The Chloride concentrations in the period of sampling for the upstream station B (Control) are 12.912 mg/l. And the two downstream treatment stations: Station A 26.378 mg/l, and station C 32 mg/l.

#### Gasterosteus aculeatus

Linnaeus, 1758 individuals were sampled using electrofishing equipment. The field sampling permit was obtained from the fisheries department of the state of Nordrhein-Westfalen, Germany. Fish were then quickly euthanized using an overdose of tricaine methanesulfonate (MS222) before harvesting the gill tissue, immediately fixing it in RNAprotect (QIAGEN), and storing it at - 20°C. In total nine samples were obtained, three from the control (Station B), three from the mid-salinity station (Station A), and three from the high-salinity station (Station C).

### RNA isolation and Illumina sequencing

The fixed tissue of each sample was homogenized in a solution of buffer 700μl RLT with a 10μl:1ml concentration of β-mercaptoethanol in a FastPrep-24™ (MP Biomedicals) bead beater during 30s at 5 m/s in a screw cap vial with 1mm diameter glass beads. Afterwards, the RNA was extracted using a RNeasy Mini Kit (QIAGEN) following the manufacturer’s instructions. The quality of the extractions was checked using a Nanodrop 1000 Spectrophotometer (Peqlab Biotechnologie, Germany), ensuring that the concentration was >50ng/μl, and that OD 260/280=1.8-2.1, and OD 260/280>1.5. Next, the RIN^e^ value quality was assessed using the RNA ScreenTape system in an Agilent 2200 TapeStation, ensuring that the samples had a RIN^e^ score over 7.0. An Illumina Tru-Seq™ RNA Sample Preparation Kit (Illumina, San Diego, CA, USA) was applied in generating sequencing libraries following the manufacturer’s protocol. After purifying, Illumina sequencing was carried out on an Illumina/Hiseq-2500 platform (Illumina, San Diego, CA, USA) to generate 100 bp paired-end reads with 50 million reads depth by the Cologne Center for Genomics (CCG), Germany.

### Sequence quality control, de novo assembly, and annotation

Raw sequences of all nine samples were checked in FASTQC [39] for quality, followed by a quality improvement via an adaptor and low-quality reads filtering using TRIMMOMATIC [40]. TRINITY v2.9.1 [41] was used to create a de novo assembly using the nine pairs of clean sequences. The reads of the raw nine paired sequences were aligned to the de novo transcriptome using SALMON [42] as the abundance estimation method. BUSCO v5.2.2 [43] was used to assess transcriptome assembly completeness by searching against the actinopterygii_odb10 (Creation date: 2021-02-19) dataset. The expression matrix was generated and the low expressions transcripts (minimum expression of 1.0) were filtered using TRINITY v2.9.1, the expression levels were normalized as transcripts per million transcripts (TPM). Identification of likely protein-coding regions in transcripts was performed with TRANSDECODER v5.5.0 [44]. The filtered transcriptome sequencing reads were aligned to protein, signal peptides, and transmembrane domains using the tools, DIAMOND v2.0.8 [45], SIGNALP 6.0 [46], TMHMM v2.0 [47], and, HMMER v3.3.2 [48]. The de novo transcriptome was then functionally annotated using the tool TRINOTATE v3.2.2 [41].

### Differential gene expression analysis

Before the differential gene expression analysis, the SALMON tool was used to align the clean reads of the three sites to the filtered transcriptome and the mapping rate was calculated. The mapping tables were merged according to the stations and using TMM [49] normalization. Differential expression analysis with three biological replicates of the two stations with higher chloride (A and C) were compared to the reference condition (Station B) using the DESeq2 [50] package, the fold change cut-off was set at >2, and the FDR≤0.05. Then, a volcano plot using GGPLOT2 [51] was used to represent the differentially expressed genes (DEGs) between the treatment and control sites. The significant upregulated and downregulated DEGs ontology terms were annotated and enrichment of analysis was performed using ShinyGo [52] (version 0.61). The genes identified in the GO analysis to be related to the pathways involved in osmoregulation were extracted and listed. The assembly was performed in the High-performance computing system of the university of Cologne (CHEOPS) and the rest of the analysis was performed using the Galaxy project platform [53].

### RNA-seq qPCR validation

To validate the results from the RNA-seq analysis, a subset of eight osmoregulation-related upregulated genes was chosen and assessed via real-time quantitative PCR analysis (qPCR). The primer design was performed using the Primer-Blast software [54], and all primer sequences are listed in Table 1. Firstly, reverse transcription of the total RNA samples was performed using the ReverAid first-strand cDNA synthesis kit (Thermo Scientific), and the resulting cDNA was diluted 10-fold and used as a template for qPCR. The qPCR was performed using PowerUp™ SYBR™ Green Master Mix in a StepOne Real-Time PCR system following the manufacturer’s indications. The qPCR was carried out in triplicate for each sample with a final well volume of 20μl. An initial 10-minute hot start at 95°C, followed by 45 cycles of 30 seconds of denaturation at 95°C and 1-minute annealing at 59°C. Followed by a melting curve ranging from 60°C to 95°C with data acquisition every 0.3°C. Each gene was assessed using the 2^-△△Ct^ method, with *GADPH* as the housekeeping gene for normalization, the optimized sequence for the gene was obtained from [33]. Finally, the sample with the lower expression of the control station was used as the reference to compare the expression.

**Table 1.**
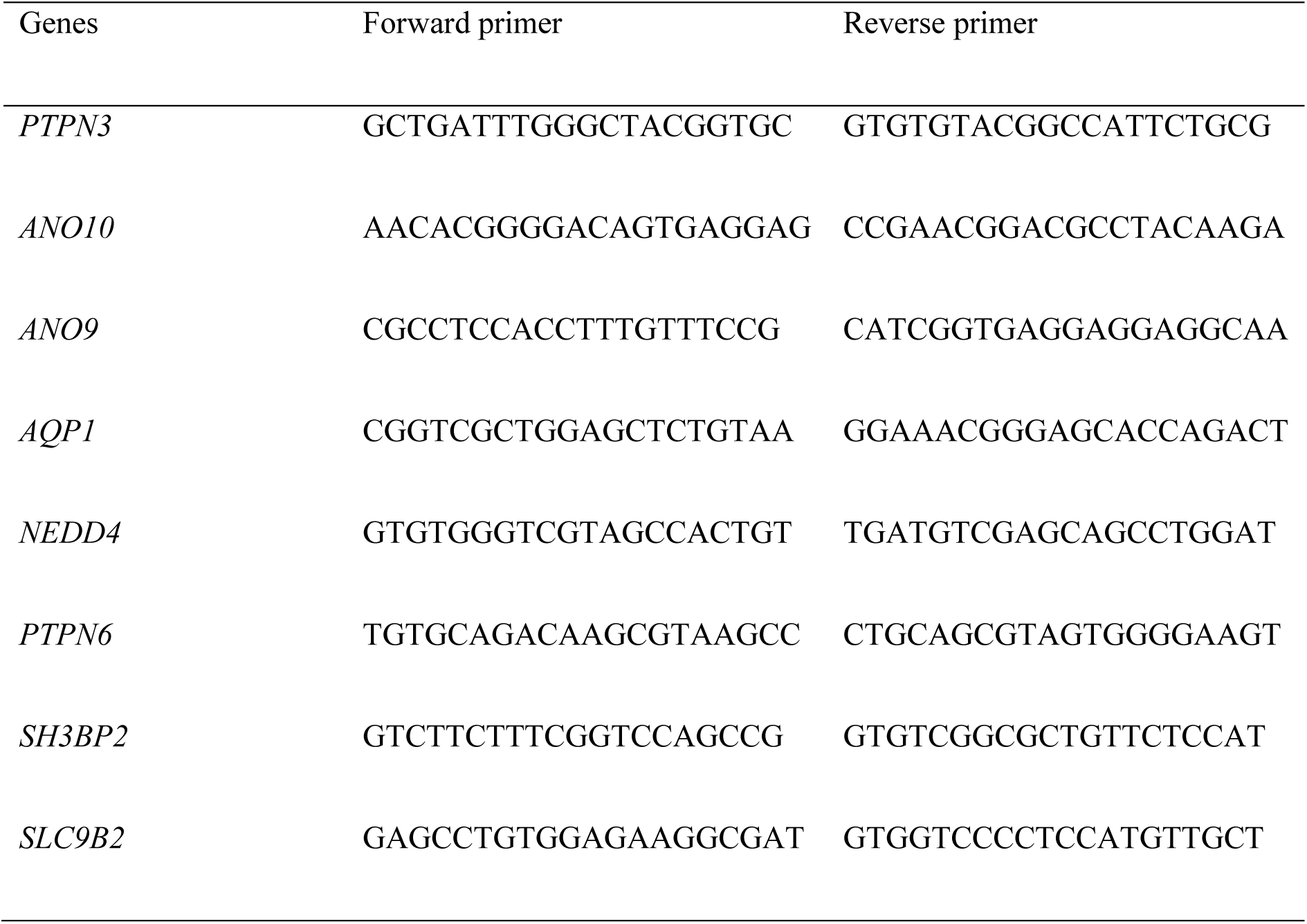
Primer sequences used for differentially expressed genes used for qPCR validation.

## Results

### Sequence quality control, de novo assembly, and annotation

The RNA-seq was performed on the library products from the gill of *G. aculeatus*, resulting in an average of 47,945,147, 64,032,899.33, and 68,881,883.67 total reads from stations A, B and C respectively. The quality score %Q30, which states the sum of the reads with a base call accuracy of 99, 9% was 99.32%, 99.36% and 99.37% respectively. After quality control, an average of 44,571,717, 59,441,980.67 and 64,633,936 were retained for stations A, B and C respectively. All the retained reads had >Q30. The assembly with all sequences produced a raw transcriptome of 559.3mb, with a N50 of 3797bp, with 3319 complete BUSCO’s and a completeness of 91.2%. The transcriptome was filtered for low-expression transcripts, and from a total 340418 in the raw transcriptome, 98643 were retained (28.98%). The filtered transcriptome outcome was 182.4mb of sequences, with a N50 of 3735bp, 2334 complete BUSCO’s and a completeness of 64.1%. In total 50155 transcripts were annotated which represents 50.8% of the de novo transcriptome.

### Differential gene expression analysis

The mapping rate of the individual sequences to the filtered transcriptome was an average of 60.68%, 59.78% and 60.18% on stations A, B and C respectively. When comparing the station with mid-salinity (A) with the reference site (B), a total of 627 genes (306 down- and 321-regulated) (Figure 2). For the comparison of the station with high-salinity (C) with the reference site (B), a total of 611 genes (270 down- and 341regulated) (Figure 3). In both heatmaps the pattern of expression was similar in all three sample replicates per station. The significant DEGs were assigned to three major Gene Ontology (GO) categories: biological process, cellular component, and molecular function. The DEGs were classified in a total of 200 and 193 higher-level GO terms for A and C, respectively, relative to the reference station (S1 File). Our comparative analysis of the high-level GO terms within the significant DEG dataset, identified potential osmoregulation-related terms, including transporter activity (GO:0005215), transmembrane transporter activity (GO:0022857), and channel regulator activity (GO:0016247).

**Figure 2.**
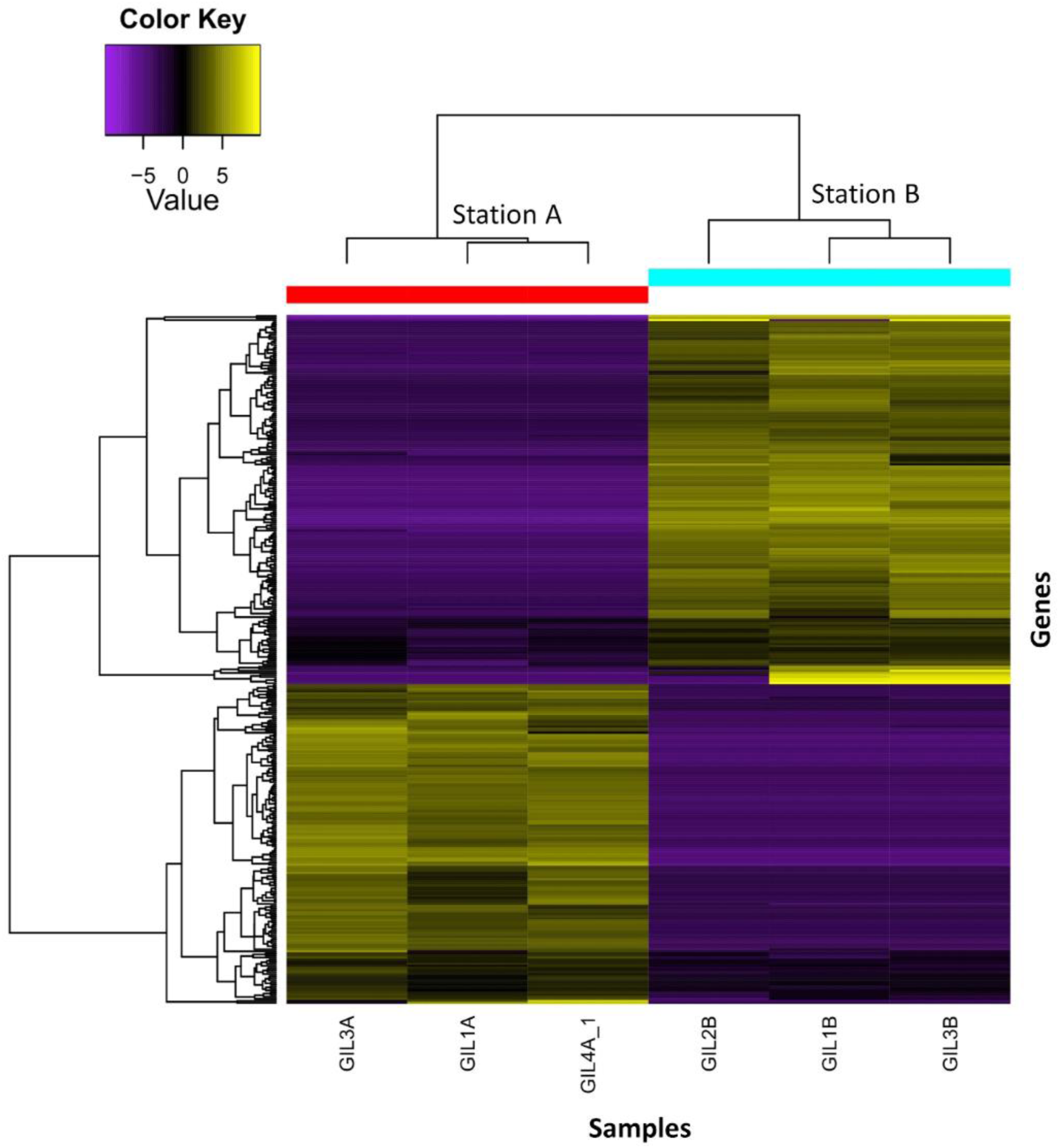
Heatmap displaying 627 differentially expressed genes between the mid-salinity station (A) and the reference station (B). The X-axis represents sample replicates per station, and the Y-axis represents individual gene expression. Upregulated genes are shown in yellow, with brighter colors indicating higher expression values. In contrast, purple shades indicate downregulated genes, with the brightest shade indicating the strongest downregulation.

**Figure 3.**
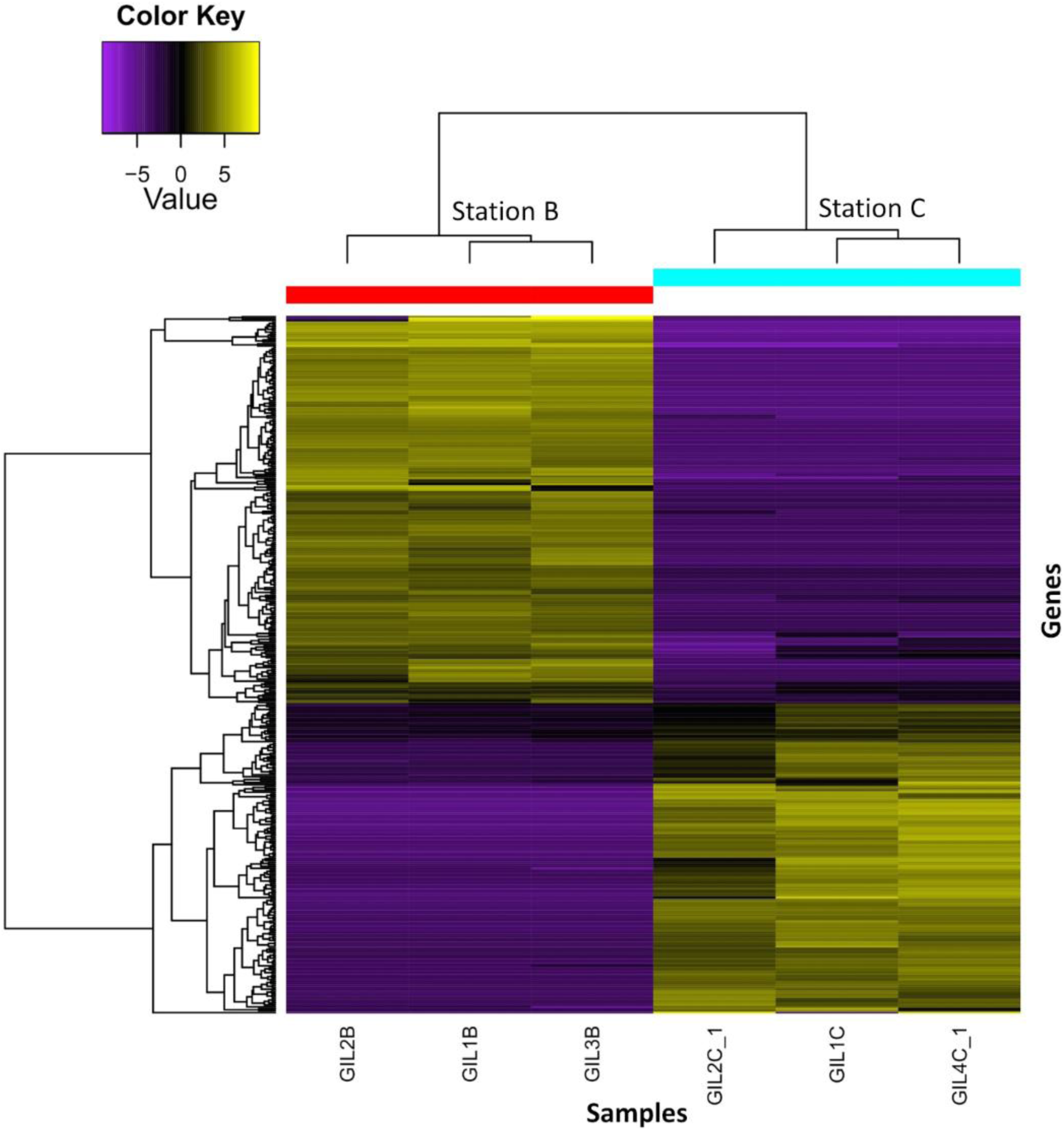
Heatmap displaying 611 differentially expressed genes between the high-salinity station (A) and the reference station (B). The X-axis represents sample replicates per station, and the Y-axis represents individual gene expression. Upregulated genes are shown in yellow, with brighter colors indicating higher expression values. In contrast, purple shades indicate downregulated genes, with the brightest shade indicating the strongest downregulation.

The GO ontology analysis was used to find which terms were enriched in the two higher salinity stations compared with the reference site for the upregulated and downregulated genes separately. Furthermore, to identify the candidate DEGs related to these particular osmoregulation-related GO terms the gene codes were extracted and assigned to the correspondent functional category term (S2 File). For the upregulated genes we found a set of enriched GO terms for the three categories, molecular function, cellular component and biological process (Figure *4*). The main enriched GO terms related to osmoregulation in the upregulated DEGs in the biological process GO category for station A vs B are: Regulation of sodium ion transmembrane transport (GO:1902305 and GO:1902306), regulation of sodium ion transmembrane transporter activity (GO:2000649 and GO:2000650), and regulation of sodium ion transport (GO:0002028 and GO:0010766). For the molecular function the GO categories main enriched terms were related with the activation and regulation of sodium channels (GO:0017080, GO:0019871, GO:0016248, GO:0016247). The main enriched GO terms related to osmoregulation in the upregulated DEGs in the biological process GO category for station C vs B are: Regulation of sodium ion transport (GO:0002028), regulation of ion transmembrane transporter activity (GO:0032412), regulation of transmembrane transporter activity (GO:0022898), regulation of transporter activity (GO:0032409), regulation of ion transmembrane transport (GO:0034765) and sodium ion transmembrane transport (GO:0035725). For the molecular function GO category in C vs B the osmoregulation-related enriched term was: Sodium channel regulator activity (GO:0017080). For the cellular component category only the osmoregulation-related term transport vesicle (GO:0030133) was found to be enriched.

**Figure 4.**
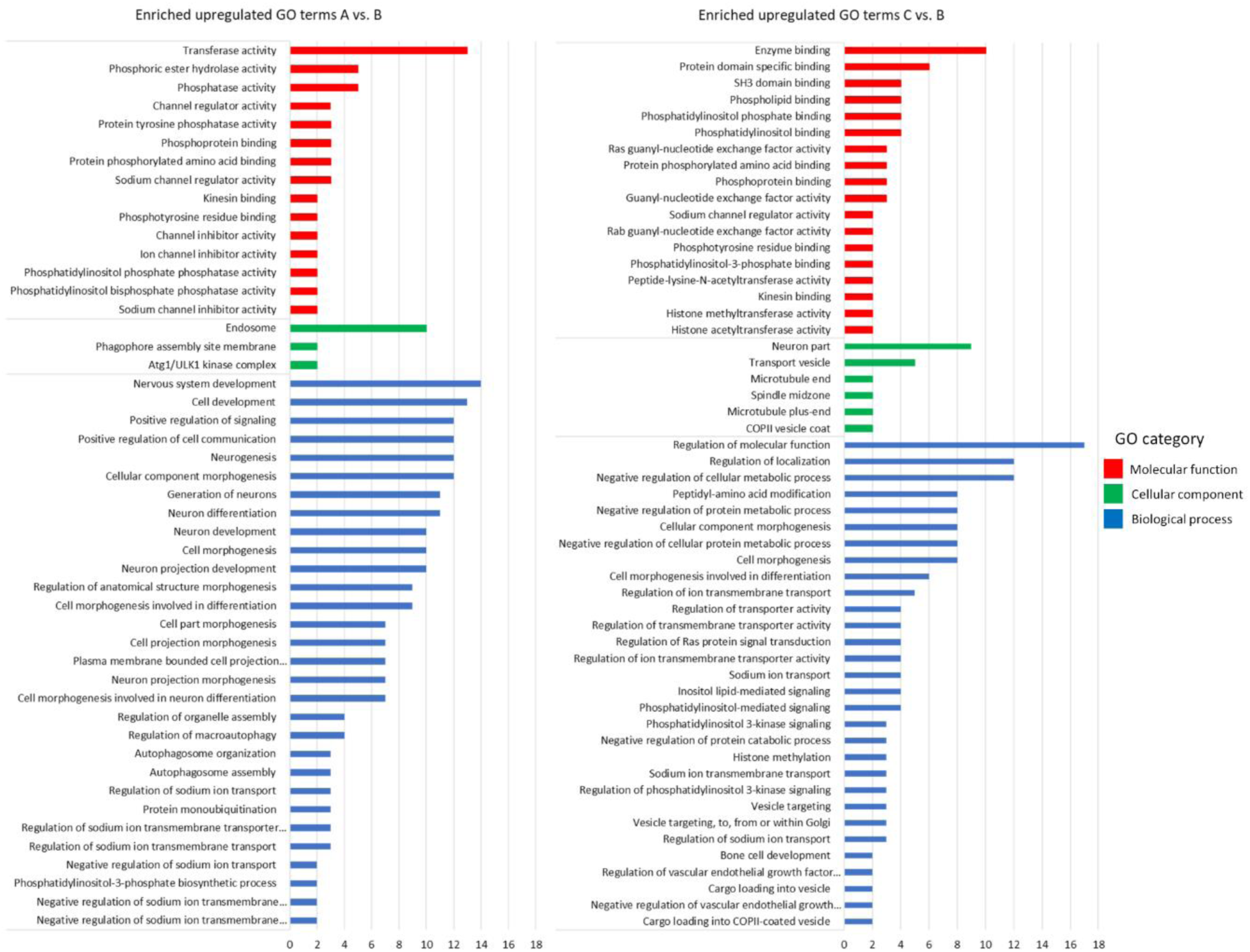
Gene ontology analysis of enriched upregulated genes for stations A and C compared to the reference site B. The number of genes is displayed on the X axis, while the Y axis represents GO categories colour-coded based on molecular function (red), cellular component (green), and biological process (blue). Could you unify the X axis to make a comparison easier? (max value AB= 16, max value CB = 18)

For the downregulated genes we found a set of enriched GO terms for the three categories, molecular function, cellular component and biological process in the A vs B and only in the molecular function and biological process terms for C vs B (Figure 5). For station A vs B, we found enriched GO terms related to osmoregulation only in the molecular function GO category: Anion binding (GO:0043168) and ATP binding (GO:0005524). The main enriched GO terms related to osmoregulation in the downregulated DEGs are in the molecular function GO category for station C vs B and are mostly related to the Ras, Rho and Rab small GTPases terms. These related terms are: Ras guanyl-nucleotide exchange factor activity (GO:0005085), Regulation of Ras protein signal transduction (GO:0046578), Rho guanyl-nucleotide exchange factor activity (GO:0005085), Rho GTPase binding (GO:0017048), GTPase regulator activity (GO:0030695), GTPase activator activity (GO:0005096) and Rho GTPase binding (GO:0031267).

**Figure 5.**
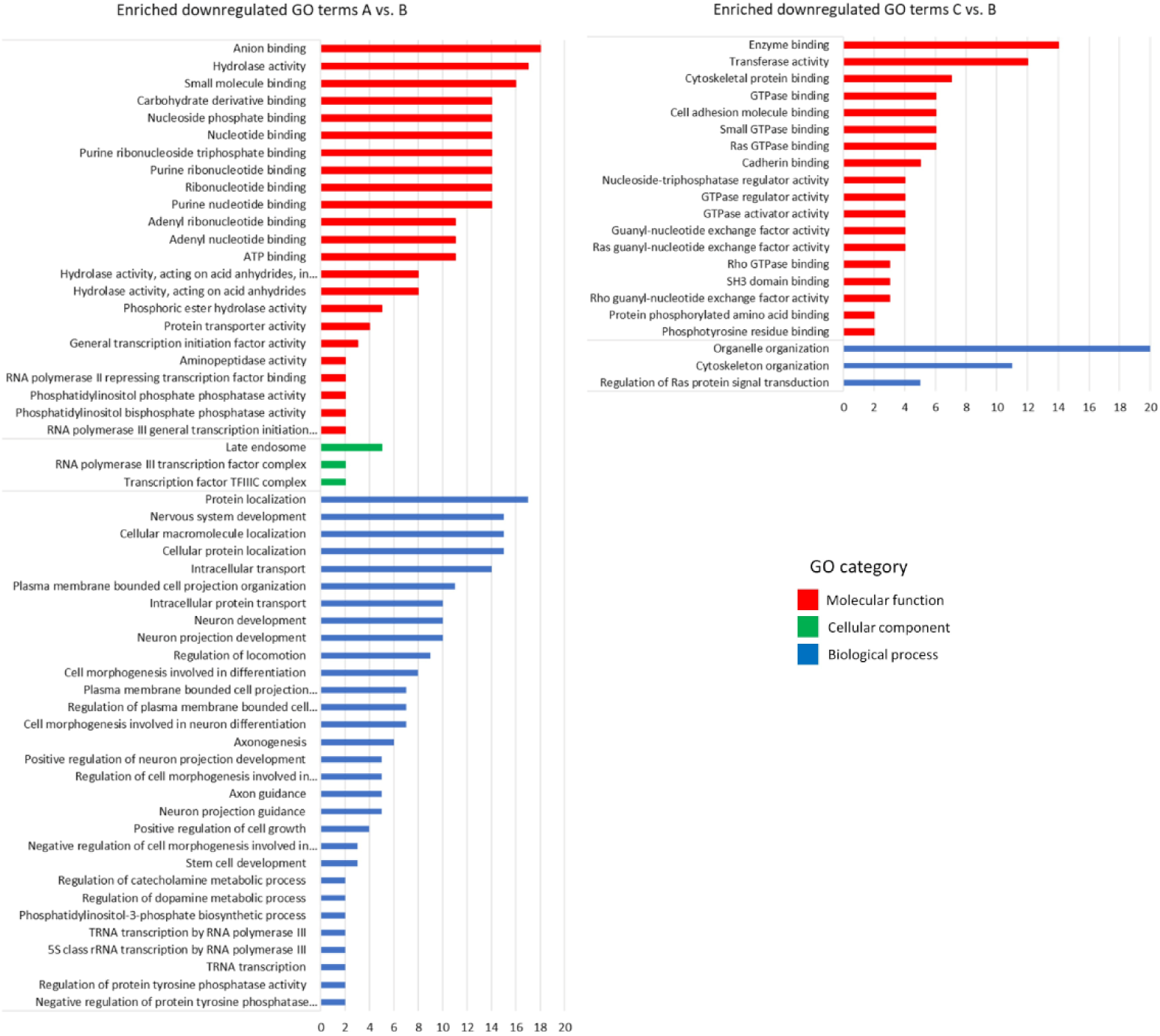
Gene ontology analysis of enriched downregulated genes for stations A and C compared to the reference site B. The number of genes is displayed on the X axis, while the Y axis represents GO categories colour-coded based on molecular function (red), cellular component (green), and biological process (blue). X axes – see above

From the enriched GO terms that were identified to be related to osmoregulation for both higher salinity stations, we extracted all the DEGs and listed them in S3 File. Then we performed a literature search of all the extracted genes and listed the DEGs with referenced osmoregulation involvement (Table *2*). For station C vs B, the genes with the higher upregulation were the protein transporters *SEC31A*, followed by *SEC24B*. Then, the regulators of the ion channel and ion transporters *PTPN3*, *PTPN6*, *AHCYL1* and *NEDD4* were also found to be upregulated. In the last position were other genes involved in ion transport, the cation channel protein anoctamin 9 *ANO9* and the solute carrier *SLC9B2*. The strongest downregulated genes in C vs B, were the Ras, Rho and Rab small GTPases related genes *ITPKB* and *AKAP13*. And finally, the genes involved in ion transport, the ion channel *TRPM7* and the solute carrier *SLC9B2*. For station A vs B, the genes with the higher upregulation were the regulators of ion channel and ion transporters *PTPN3*, *NEDD4* and *NEDD4L*. Followed closely by the channel protein-encoding gene *AQP1* and the solute carrier *SLC35B2*. In the last position with lower expression were the channel regulators *PRF1*, *CLCN7*, the solute carrier *SLC6A15*, and the channel protein Anoctamin 10 *ANO10.* The strongest downregulation was recorded for the membrane tension regulator *SCARB2*, the transporter *ABCE1*, the channel regulators *VPS35* and *VP26C* and the protein complex gene *PIKFYVE*. In the last position was the anion exchanger *ATAD2*.

**Table 2.** List of the selected candidate genes involved in osmoregulation generated from the Gene Ontology analysis. Upregulated genes are highlighted in red, while downregulated genes are highlighted in blue, with intensity indicating expression level (Log2 FC). The false discovery rate (FDR) value is also included, indicating that an FDR p<0.05 suggests a non-random differential expression of the gene.

| Group | Gene | Description | Log2 FC | FDR(p) |
| --- | --- | --- | --- | --- |
| Station C vs. Station B | *SEC31A* | Protein transport protein Sec31A, ABP125, ABP130, SEC31-like protein 1, SEC31-related protein A, Web1-like protein | 12.35 | 6.54E-17 |
|  | *SEC24B* | Protein transport protein Sec24B, SEC24-related protein B | 11.71 | 4.12E-16 |
|  | *PTPN3* | Protein tyrosine phosphatase non-receptor type 3 | 11.17 | 2.79E-11 |
|  | *AHCYL1* | Adenosylhomocysteinase like 1 | 10.57 | 8.07E-06 |
|  | *NEDD4* | Nedd4 e3 ubiquitin protein ligase | 10.06 | 3.00E-09 |
|  | *PTPN6* | Protein tyrosine phosphatase non-receptor type 6 | 10.00 | 7.48E-14 |
|  | *ANO9* | Anoctamin 9 | 9.47 | 1.63E-07 |
|  | *SLC9B2* | Solute carrier family 9 member b2 | 9.15 | 6.22E-08 |
|  | *TRPM7* | Transient receptor potential cation channel subfamily M member 7 | -2.74 | 4.53E-03 |
|  | *SLCO5A1* | Solute carrier organic anion transporter family member 5A1 | -9.05 | 1.07E-10 |
|  | *AKAP13* | A-kinase anchor protein 13, AKAP-13Non-oncogenic Rho GTPase-specific GTP exchange factor | -12.53 | 3.02E-18 |
|  | *ITPKB* | Inositol-trisphosphate 3-kinase B, 2.7.1.127, Inositol 1,4,5-trisphosphate 3-kinase B, IP3 3-kinase B, IP3K B, InsP 3-kinase B | -13.11 | 3.00E-24 |
| Station A vs.<br>Station B | *PTPN3* | Protein tyrosine phosphatase non-receptor type 3 | 11.22 | 1.35E-17 |
|  | *NEDD4* | Nedd4 e3 ubiquitin protein ligase | 10.87 | 7.59E-06 |
|  | *NEDD4L* | Nedd4 like e3 ubiquitin protein ligase | 10.31 | 9.20E-11 |
|  | *AQP1* | Aquaporin 1 (colton blood group) | 8.09 | 5.37E-07 |
|  | *SLC35B2* | Solute carrier family 35 member b2 | 7.55 | 0.01 |
|  | *PRF1* | Perforin 1 | 3.65 | 0.01 |
|  | *CLCN7* | Chloride voltage-gated channel 7 | 2.48 | 6.49E-05 |
|  | *SLC6A15* | Solute carrier family 6 member 15 | 2.40 | 0.01 |
|  | *ANO10* | Anoctamin 10 | 2.23 | 0.01 |
|  | *ATAD2* | ATPase family AAA domain-containing protein 2, 3.6.1.- | -2.19 | 0.0002 |
|  | *VP26C* | Vacuolar protein sorting-associated protein 26C | -8.67 | 2.3E-04 |
|  | *PIKFYVE* | 1-phosphatidylinositol 3-phosphate 5-kinase, Phosphatidylinositol 3-phosphate 5-kinase, 2.7.1.150 | -10.39 | 3.76E-11 |
|  | *VPS35* | Vacuolar protein sorting-associated protein 35 | -10.52 | 2.24E-13 |
|  | *ABCE1* | ATP-binding cassette sub-family E member 1, 2'-5'-oligoadenylate-binding protein | -11.50 | 4.86E-11 |
|  | *SCARB2* | Lysosome membrane protein 2 | -12.67 | 3.7E-20 |

### RNA-seq data validation by qPCR

The analysis of the melting curve of the qPCR resulted in a single product for all genes. The visual comparison of the relative fold changes of the subset of eight DEGs showed a consistently positive direction in both qPCR and RNA-Seq analysis (Figure 6), which suggests that the RNA-seq analysis is reliable.

**Figure 6.**
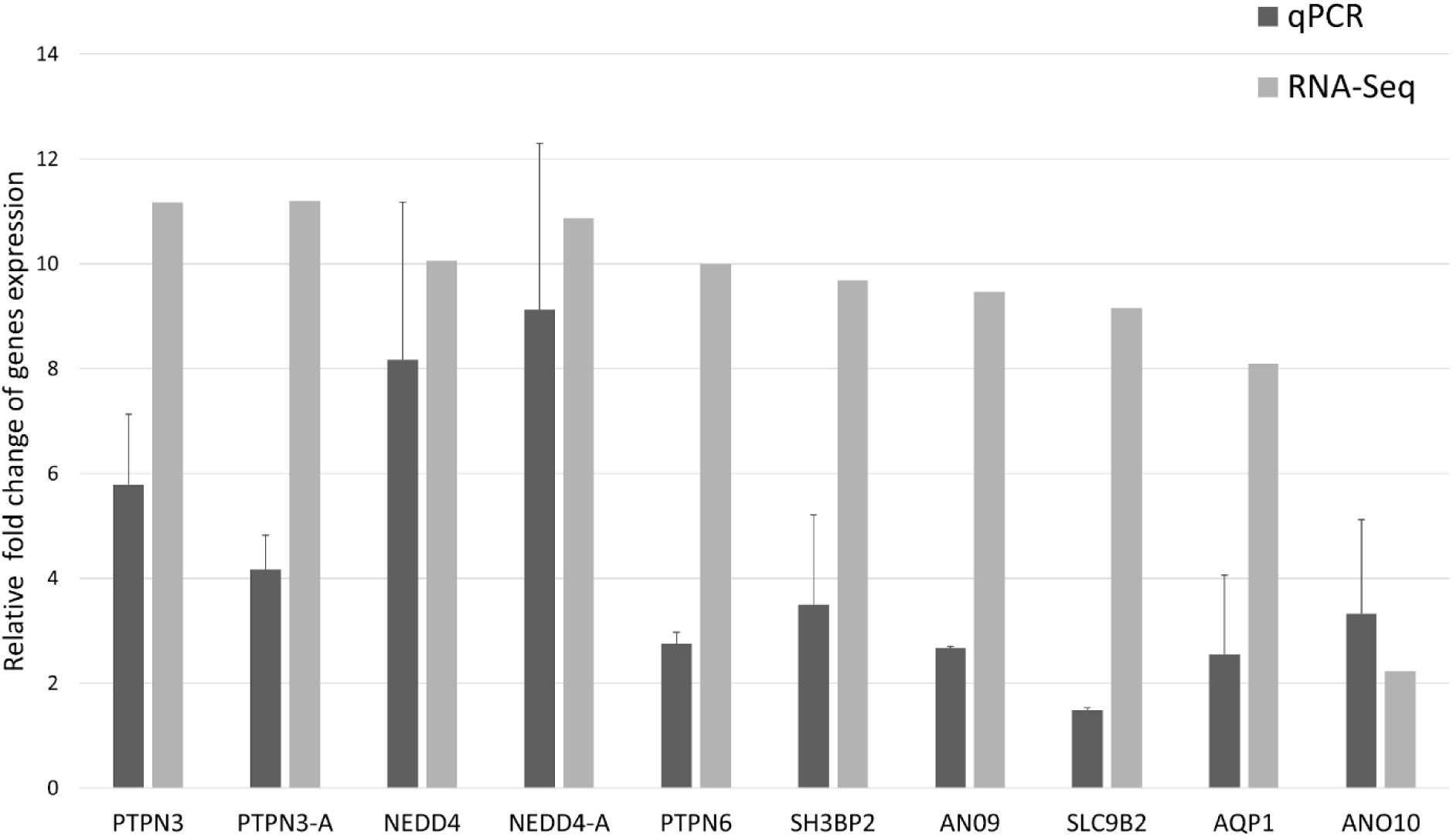
qPCR validation plot of RNA-Seq data. It shows the expression of eight genes, including *NEDD4* and *PTPN3*, in two treatment groups. *NEDD4* and *PTPN3* refers to group C vs. B, while *NEDD4*-A and *PTPN3*-A refer to group A vs. C. The gene names are on the X axis, and the Y axis displays the relative fold change in gene expression (Log2FCs). Mean ± standard deviation Log2FC values are presented.

## Discussion

Our transcriptomics analysis yielded over 50 million reads with 100% of reads surpassing Q30, and a N50 length of 3735bp, indicating high-quality raw data. The mapping rate, which was consistent with previous studies [25,55], was close to 60% for all stations, demonstrating the effectiveness and reliability of the high-throughput sequencing transcriptomic analysis. Additionally, the maintained trend in gene expression between qRT-PCR and RNA-seq validated our results and confirms their suitability for further analysis. According to our prediction, we found a differential expression of the genes in the gill tissue of *G. aculeatus* under a gradient of salinity in the field. Notably, even when chloride concentrations were well below reported toxic thresholds, our comparative transcriptomics approach was able to evidence the activation of the energetically costly osmoregulatory system. Furthermore, we were able to identify candidate genes related to increased osmoregulatory activity even for subtle changes of chloride concentration. In the ontology analysis, we found an enrichment of DEGs genes predominantly related to transmembrane transport and regulation of transport. It is to highlight, that in neither of the treatments we found DEGs with functional categories that could be related to the handling or euthanasia of the fish. The pathways and genes that we found fall in one or more of the big three classes of proteins in charge of moving ions across membranes, ion channels, ion pumps and ion transporters which are essential for all living organisms [56]. These key groups of proteins maintain the osmotic pressure in exchange for considerable quantities of energy, and together receive the name “Transportome” [57]. The transportome is in charge of maintaining biological homeostasis through a balance between the intracellular and extracellular environments and currently accounts for 10% of the protein-coding human genome [58]. For instance, studies with Chinese mitten crab, Guppy *Poecilia reticulata*, *Acipenser baeri*, hybrid tilapia, and *G. aculeatus* showed a significant number of DEGs involved in ion transport and transmembrane transport [22,24,25,28,59]. These results confirm the importance of the transportome in the adaptive response to salinity changes and align with the enrichment of genes in related categories in our study. In the following sections, we discuss the relevance of the enriched pathways for osmoregulation, the specific function of the candidate genes, and the ecological implications of our findings for euryhaline fish such as *G. aculeatus* in the face of global freshwater salinization.

### Membrane transporters DEGs

Membrane transporters were the most abundant group of candidate genes that were differentially expressed in the stations with higher chloride concentrations. The candidate genes related to these pathways are *VPS35*, *VPS26*, *SLC35B2*, *SLC9B2*, *SLCO5A1*, *SLC6A1*, *SEC31A*, and *SEC24*. The Vacuolar protein sorting-associated proteins *VPS35* and *VPS26* are components of the cargo recognition sub-complex of the retromer, and together they recognize and bind to specific sorting signals on cargo proteins to be transported back to the Golgi apparatus [60,61]. Both proteins are crucial in cell homeostasis, by regulating the transport of particles and by controlling the water reabsorption protein Aquaporin-3 (Aquaporin-3) [62]. According to the review by Verri et al., (2012) [63], the *SLCs* transporters in teleost fish, are conformed by over 50 families with 380 known members and facilitate the transportation of nearly all soluble molecules across cellular membranes. *SLC35B2* is part of the nucleoside-sugar transporter family (*SLC35*) that is involved in the endoplasmic reticulum and Golgi of eukaryotic cells [64]. Similar to our findings, differential expression of *SLC35* family members was reported in the gills of the Chinese mitten crab in response to changes in salinity [59]. A remarkable finding was the upregulation of *SLC9B2*, as the members of the *SLC9* family play a critical role in regulating pH in cells and organelles, as well as maintaining acid-base and volume homeostasis [65]. In previous studies a high upregulation of *SLC9* genes has been recorded under salinity stress, in particular, *SLC9B2* was found to function as a Na+/H+ exchanger in an experiment using Xenopus oocytes [66]. *SLCO5A1* from the *SLC5* Sodium-glucose cotransporter family and *SLC6A15* from the SLC6 Sodium- and chloride-dependent neurotransmitter transporter family were also differentially expressed in our study and are also important transporters found in fish [63]. Furthermore, The SLCs have shown to hold high importance in generating counter-ion fluxes to maintain ion homeostasis while avoiding overtly high membrane tension [67]. In addition to observing an enrichment of SLC proteins under salinity stress, we also found an increase in proteins related to the vesicular traffic of the SEC family, such as *SEC31A* as it was also found in Malik and Kim (2021), and *SEC24B*. The impairment of the expression by glucocorticoids of members of the *SEC24* family has been related to the aberrant ion and macromolecular transport in Zebrafish [68]. Overall, the findings suggest that these candidate genes play an important role in maintaining ion and macromolecular transport under salinity stress.

### Ion channel regulation and transport DEGs

Another important group of DEGs enriched terms was the one related to ion channel regulation and transport. Examples of this that we pinpointed as candidate genes are *PTPN3*, *PTPN6*, *NEDD4*, *NEDD4L*, *AHCYL1*, *AQP1*, *TRPM7*, *PRF1*, *CLCN7*, *ANO9* and *ANO10*. This holds particular importance as ion channels such as voltage-gated, ligand-gated, and second messenger-gated channels, nonselective cation channels, and epithelial Na+ and Cl-channels are key to maintaining cellular homeostasis [69]. And as described in Davis et al., (2001) [70] protein tyrosine phosphatases as *PTPN3* and *PTPN6* are involved in tyrosine phosphorylation, which is the primary means of regulating the majority of voltage-gated, ligand-gated, and second messenger-gated channels. *PTPN3* has been found to regulate the high-osmolarity glycerol (HOG) *MAPK* pathway, which is essential for yeast survival in a hyperosmotic environment [71]. Other important regulators that we propose as candidate genes are *NEDD4* and *NEDD4L*, which have been found to mediate ubiquitination and degradation of *AQP2* which is a key protein in water homeostasis [72]. Also, the *AHCYL1* protein has been identified as a regulator of the intracellular Ca2+ channel inositol 1,4,5-trisphosphate (IP3), and multiple ion channel and ion transporters [56]. *AQP1* is a member of the aquaporin gene family, a group of channel proteins that are involved in the transport and other solutes in the presence of an osmotic gradient. *AQP1* differential expression has been previously detected in fish gill, kidney and gut tissue in response to salinity stress for various species [73,74]. *TRPM7* is a plasma-membrane protein expressed ubiquitously, which belongs to the melastatin-related transient receptor-potential ion channel *TRPM* subfamily and possesses both ion channel and α-kinase domains. It has been observed that *TRPM7* is involved in channel regulation via changes in intracellular Ca2+ concentration when subject to osmolality gradients [75]. The perforin (*PRF1*) was also differentially expressed, which is relevant as it is known to be able to polymerize and form channels in cell membranes of target cells [76]. Also relevant was the differential expression of the chloride channel (*CLC7*), which was also found to be expressed under osmotic stress in the gills of tilapia under osmotic stress [28]. And finally we found upregulation of both transmembrane proteins anoctamin 9 and 10 (*ANO9* and *ANO10*). Anoctamins are a family of Ca2+-activated Cl− channels and phospholipid scramblases that have been shown to support cell volume regulation [77,78]. The differential expression of these genes highlights their potential role in cellular homeostasis regulation under stress conditions. Further studies on these candidate genes could lead to a better understanding of the adaptive mechanisms used by fish to cope with environmental stressors.

### Small GTPases, anion and ATP binding DEGs

Another set of pathways that were found to be differentially expressed are those related to the Ras, Rho and Rab small GTPases, anion and ATP binding genes. The importance of these pathways lies in their direct interaction with ion channels to regulate ion homeostasis [79]. Furthermore, Ras GTPases are involved in the process of membrane thinning and curvature with effects on osmoregulation [80]. One important candidate gene found in our study was the Rho GTPase *AKAP13*, which is a key regulator of ion homeostasis under osmotic stress. In detail, *AKAP13* attracts *JIP4* and activates *NFAT5* through the Rho-type small G-proteins and p38 *MAPK* signalling pathway, regulating intracellular osmolarity [81]. *ITPKB* regulates the subcellular distribution of Rasa3, a Ras GTPase-activating protein and has been observed at is differentially expressed when hyper-osmotically stressed [82,83]. In the gene ontology categories, anion and ATP binding we highlight the candidate genes *ABCE1*, *SCARB2*, *PIKFYVE* and *ATAD2*. *ABCE1* is a member of the ABC gene family, which constitutes the largest group of transmembrane transporter proteins encoded in the human genome. These proteins utilize ATP as an energy source to facilitate the transportation of various molecules across cellular membranes keeping cell homeostasis [84]. The lysosomal integral membrane protein 2 (*SCARB2*) is a transporter of cholesterol that is suspected to control membrane tension and ion trafficking via the direct insertion of lipids [85]. *PIKFYVE* is the sole source of all the Phosphoinositide lipids PI(5)P pool in yeast membranes and is a key regulator of ion transport, and it is upregulated under hyperosmotic shock in yeast [86]. Finally, another interesting candidate is *ATAD2*, a gene from the ATPase family. Its relevance resides in the fact that ATPases are enzymes that utilize ATP to power many cellular processes including transmembrane transport and osmoregulation in fish [87,88]. Note that the differential expression of *ABCE1* and *ATAD2* provides direct evidence of the energetic cost of osmoregulation, even at low salinity concentrations in our study. The proposed candidate genes play important roles in transmembrane transport, ion trafficking, and osmoregulation.

### Ecological implications

Our study demonstrates that the gill osmoregulatory mechanism of *G. aculeatus* is activated at sublethal concentrations of chloride, and we have identified related candidate genes that could serve as indicators of salinity stress. Using a transcriptomic approach, we detected activation of osmoregulatory pathways in both study sites, even below the thresholds established by German authorities for running waters. This finding is significant, as transcriptomics has been successful in determining the sublethal effects of various stressors and giving new insights into the management of species and ecosystems. For example, Komoroske et al., (2016) [35], were able to determine that the endangered delta smelt (*Hypomesus transpacificus*) osmoregulatory system was activated at a high energetically cost under sublethal salinity thresholds. And for the same species, again using transcriptomic coupled with physiological methods, it was demonstrated that sublethal critical effects are observed 4-6 °C below the previously established acute tolerance limits [89]. The relevance of this transcriptomic study was further demonstrated when Brown et al., (2016) [90], combined the information on these sublethal thresholds to model the future habitat suitability of this species. Based on these precedents, we predict that our results have the potential to inform models on the expanding distribution beyond its native range of *G. aculeatus* in response to freshwater salinization. *G. acualeatus* populations have been increasing rapidly in several ecosystems around the world in response to altered conditions, with important negative impacts on native species that have led to consider it an invasive species in places such as lake Constance (Germany) [91,92]. An effect that has been previously described for other invasive species that have demonstrated adaptive advantage in increasing salinity scenarios, increasing their populations and their distributions [93,94].

It is worth noting that the candidate genes related to osmoregulation discussed in our study, which showed differential expression, have primarily been identified in manipulative experiments involving significant changes in salinity. This highlights the relevance of our findings, as we have identified the same candidate genes within the subtle salinity gradient of our field experiment, suggesting their potential as biomarkers for rapidly assessing sublethal salinity stress in field applications. As our understanding of the mechanisms of osmoregulation advances with the use of transcriptomics, more biomarkers or candidate genes are identified. The expression of these selected biomarkers can then be used in transcriptomic arrays to evaluate the responses of individual fish to several important biological and environmental stressors such as water-quality stressors, aquatic invasive species, and climate change [95]. Thus, further studies into the mechanisms and candidate genes that we present here for sublethal salinity stress have the potential to have an impact on the conservation and management of freshwater ecosystems in the prospect of increasing freshwater salinization.

## Conclusion

In conclusion, our transcriptomics analysis of the gill tissue of *G. aculeatus* under a gradient of salinity in the field has revealed a significant differential expression of genes related to membrane transport and regulation of transport. Our results support the notion that the transportome plays a crucial role in the adaptive response to salinity changes in euryhaline fish and highlight the activation of energetically costly osmoregulatory systems even when chloride concentrations are below the established toxic thresholds. We would like to highlight our finding of a set of candidate genes related to the sublethal concentration of chloride that is involved in transmembrane transport and ion homeostasis and can be key to the further development of transcriptomic tools for evaluating stress response. Overall, our study underscores the importance of understanding the molecular mechanisms underlying osmoregulation in the face of global freshwater salinization and provides a basis for further investigation into the impact of environmental stressors on fish populations.

## Acknowledgements

This paper benefited from the multiple discussions within the Collaborative Research Centre 1439 RESIST (Multilevel Response to Stressor Increase and Decrease in Stream Ecosystems; www.sfb-resist.de). We thank Eduardo Acosta for the interesting discussion around the manuscript and the support in laboratory technics training and documentation. And Kristin Peters for her feedback on the site location map preparation. We also thank the Regional Computing Center of the University of Cologne (RRZK) for providing support and computing time on the High Performance Computing (HPC) system CHEOPS. And finally we thank the Cologne Center for Genomics (CCG), for their technical support on sample preparation and sequencing.

## Supporting information

**S1 File. High-level gene ontology classification of the DEGs genes database.** In this database, the differentially expressed genes of treatments A and C, compared to the control, are categorized into three main Gene Ontology categories and their corresponding high-level pathway categories.

**S2 File. Database of the significantly enriched gene ontology terms of the DEGs.** Database of the complete list of enriched gene ontology terms for the different treatments, where the related genes are extracted and assigned to the correspondent gene.

**S3 File. Database of the DEGs related to osmoregulatory gene ontology terms.** In this database all the genes that were identified to be related with a osmoregulation-related GO term are listed and detailed information is given.

